## Supplementary File for "Modulation of riboflavin biosynthesis and utilization in mycobacteria"

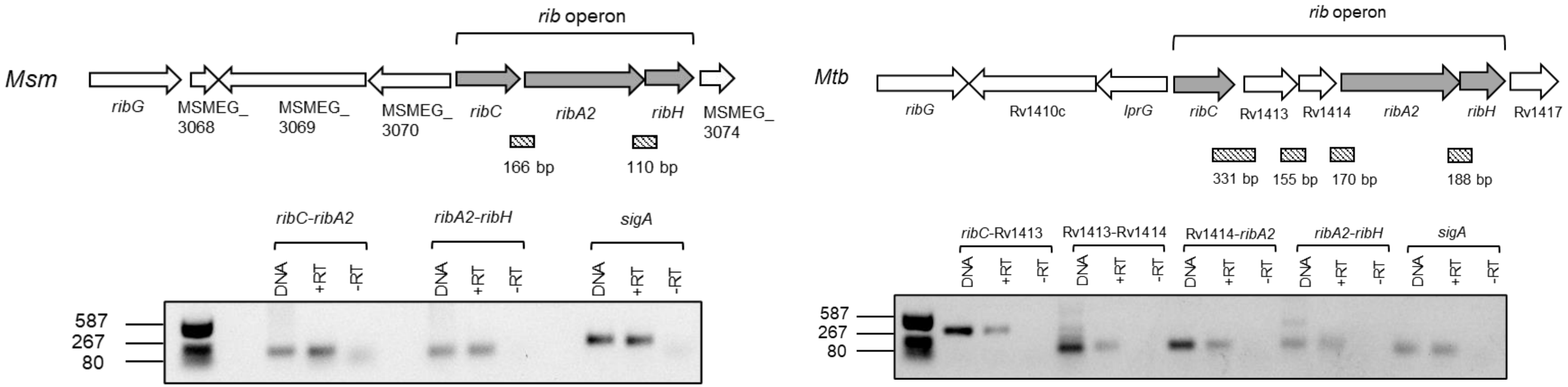

**FIG. S1.** RF biosynthesis (*rib*) operons in Msm and Mtb. Clustered *rib* genes are shown in grey. Also shown is an agarose gel electrophoresis of PCR products (sizes represented by diagonal line box) from cDNA template, using primers designed to amplify junctions between genes. Reactions were performed either in the presence (+RT) or absence of reverse transcriptase (-RT) and genomic DNA was used as a control.

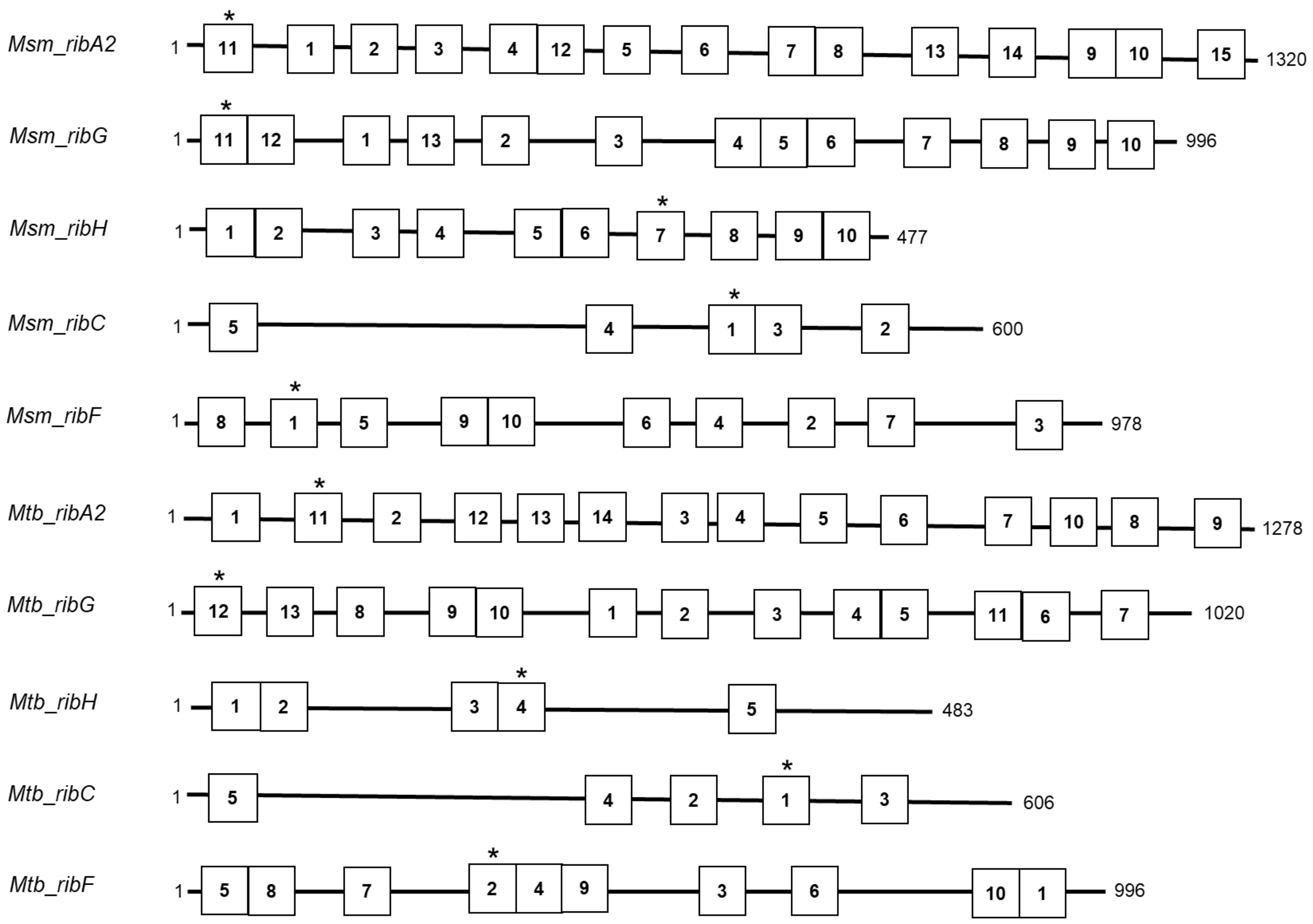

**FIG. S2.** Approximate positions of sgRNAs used to create hypomorphs in RF pathway genes. sgRNAs next to one another represent those sharing overlapping sequences. The sgRNA sequences are provided in Tables S1 and S2. Asterisks denote the sgRNA selected for detailed characterization. Not drawn to scale.

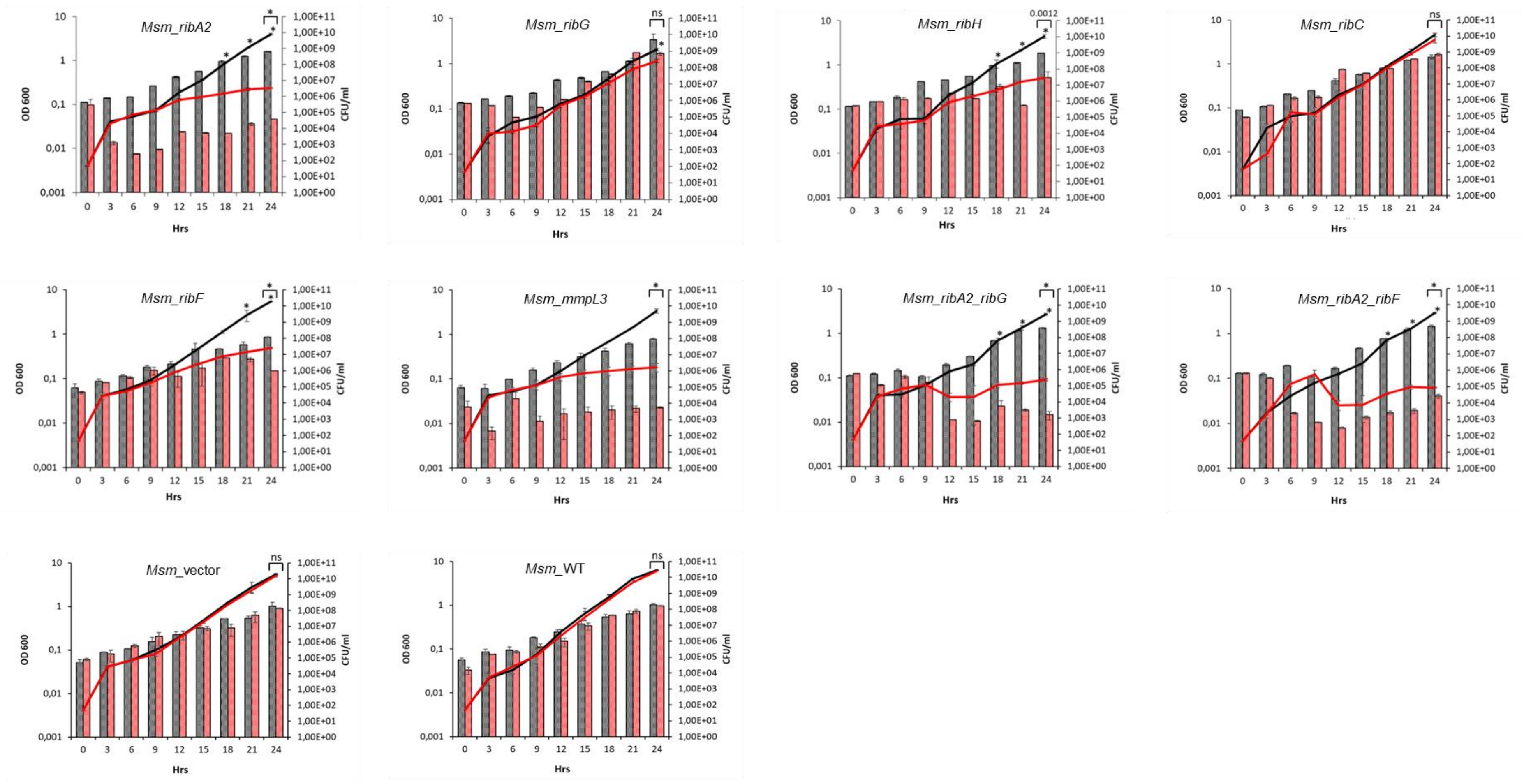

**FIG. S3**. Impact of RF pathway gene silencing on growth and viability of Msm in liquid culture. Cultures were monitored every 3 h by measuring $OD_{600}$ (solid lines) and enumerating CFUs by plating serial dilutions on 7H10 agar (columns). Cultures were grown without ATc (black solid line and black column) or in the presence of 100 ng/ml ATc (red solid line and red column). Error bars represent the SD derived from two biological replicates. Statistical comparisons were performed using a two-way ANOVA and Sidak's multiple comparison test whereby statistical significance is represented by $p < 0.0001$, denoted by an asterisk and no statistical difference represented as ns.

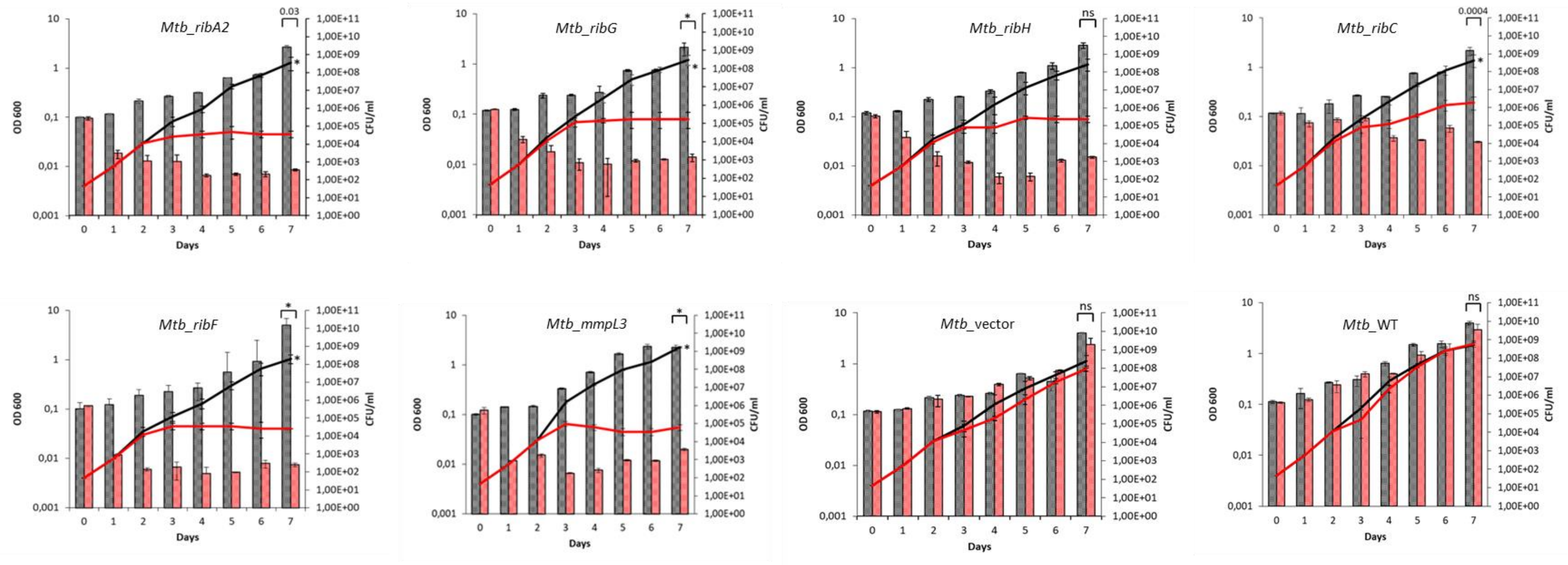

**FIG. S4**. Impact of RF pathway gene silencing on growth and viability of Mtb in liquid culture. Cultures were monitored every 24 h by measuring $OD_{600}$ (solid lines) and enumerating CFUs by plating serial dilutions on 7H10 agar (columns). Cultures were grown without ATc (black solid line and black column) or in the presence of 100 ng/ml ATc (red solid line and red column). Error bars represent the SD derived from two biological replicates. Statistical comparisons were performed using a two-way ANOVA and Sidak's multiple comparison test whereby statistical significance is represented by $p < 0.0001$, denoted by an asterisk and no statistical difference represented as ns.

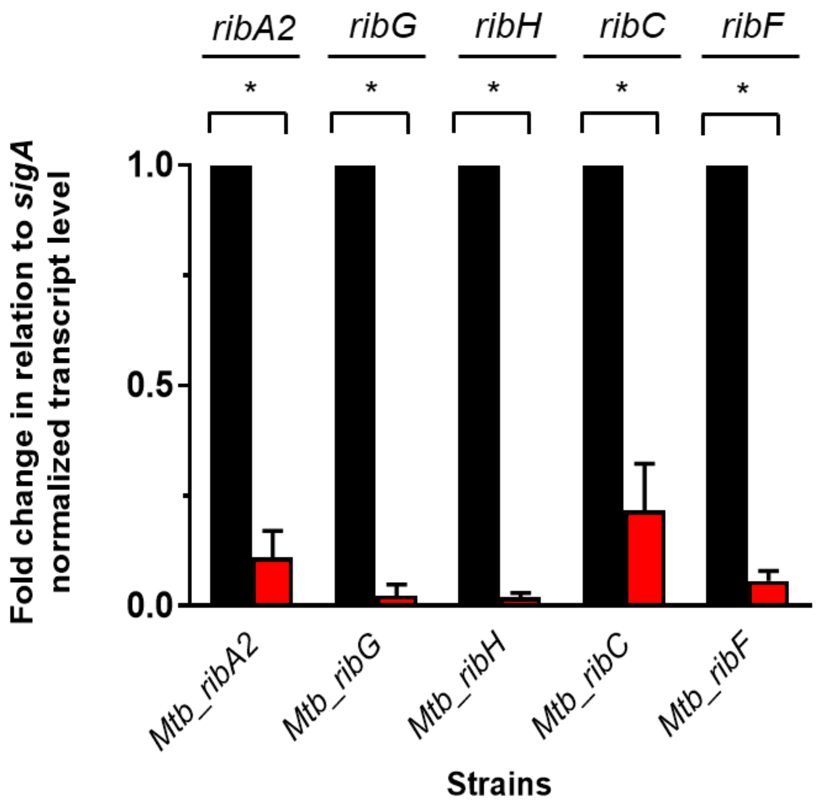

**FIG. S5**. Effective knockdown of RF pathway genes in Mtb by induced CRISPRi. The fold change in *sigA*-normalized expression of the targeted gene transcripts was determined by qRT-PCR analysis from cultures of Mtb hypomorphs grown in the absence (black) or presence of ATc (red). Error bars represent the SD derived from three biological replicates. Statistical comparisons were performed using a two-way ANOVA and Sidak's multiple comparison test whereby statistical significance is represented by $p < 0.0001$, denoted by an asterisk.

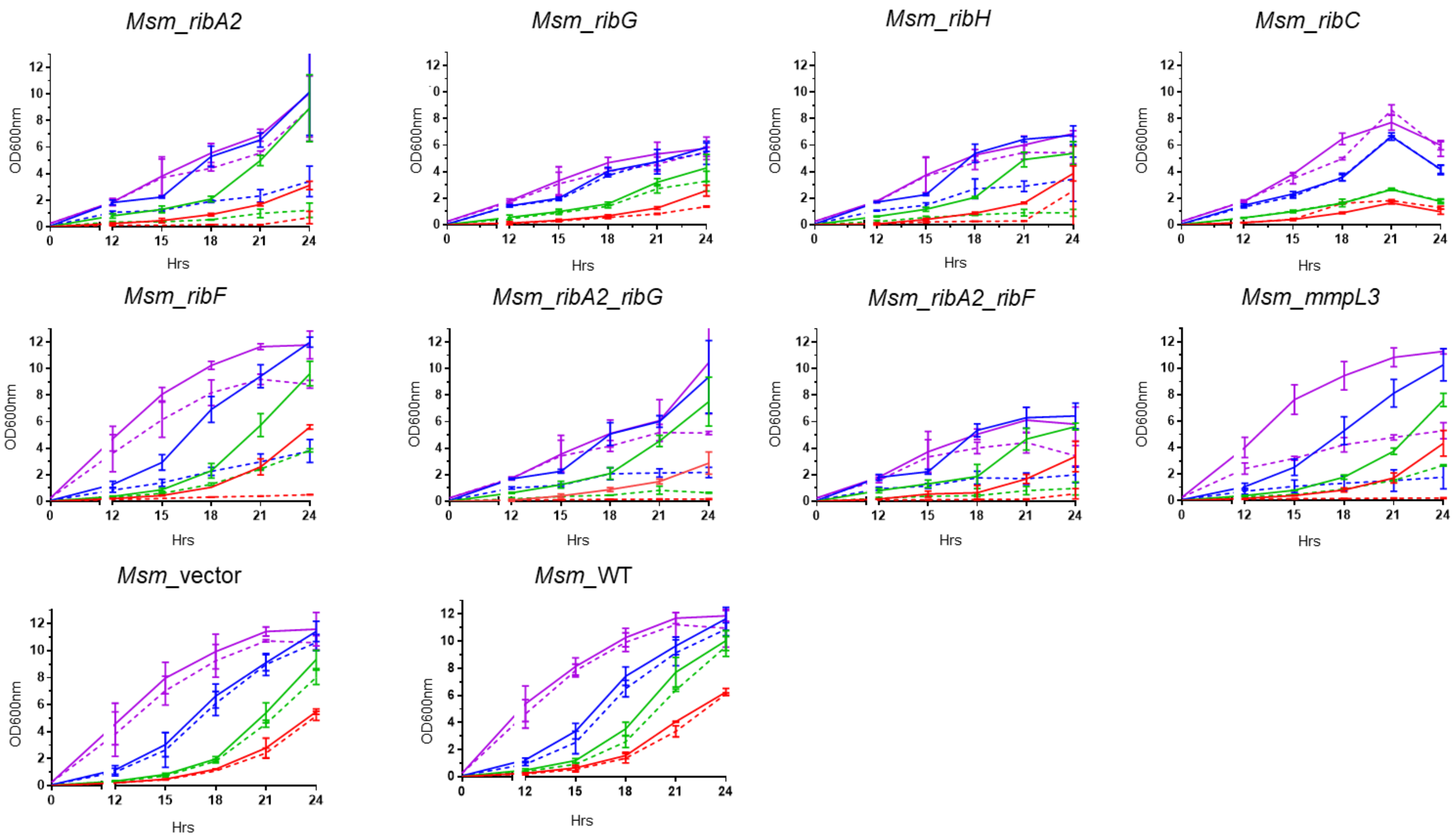

**FIG. S6**. Efficacy of CRISPRi-dependent growth repression of Msm hypomorphs as a function of inoculum size. Kinetics of growth of Msm hypomorph cultures inoculated with cells at an $OD_{600}$ value of 0.004 (red), 0.016 (green), 0.06 (blue) or 0.25 (purple). The solid line represents growth in the absence of ATc and the dashed line in the presence of 100 ng/ml ATc. Error bars represent the SD derived from three biological replicates.

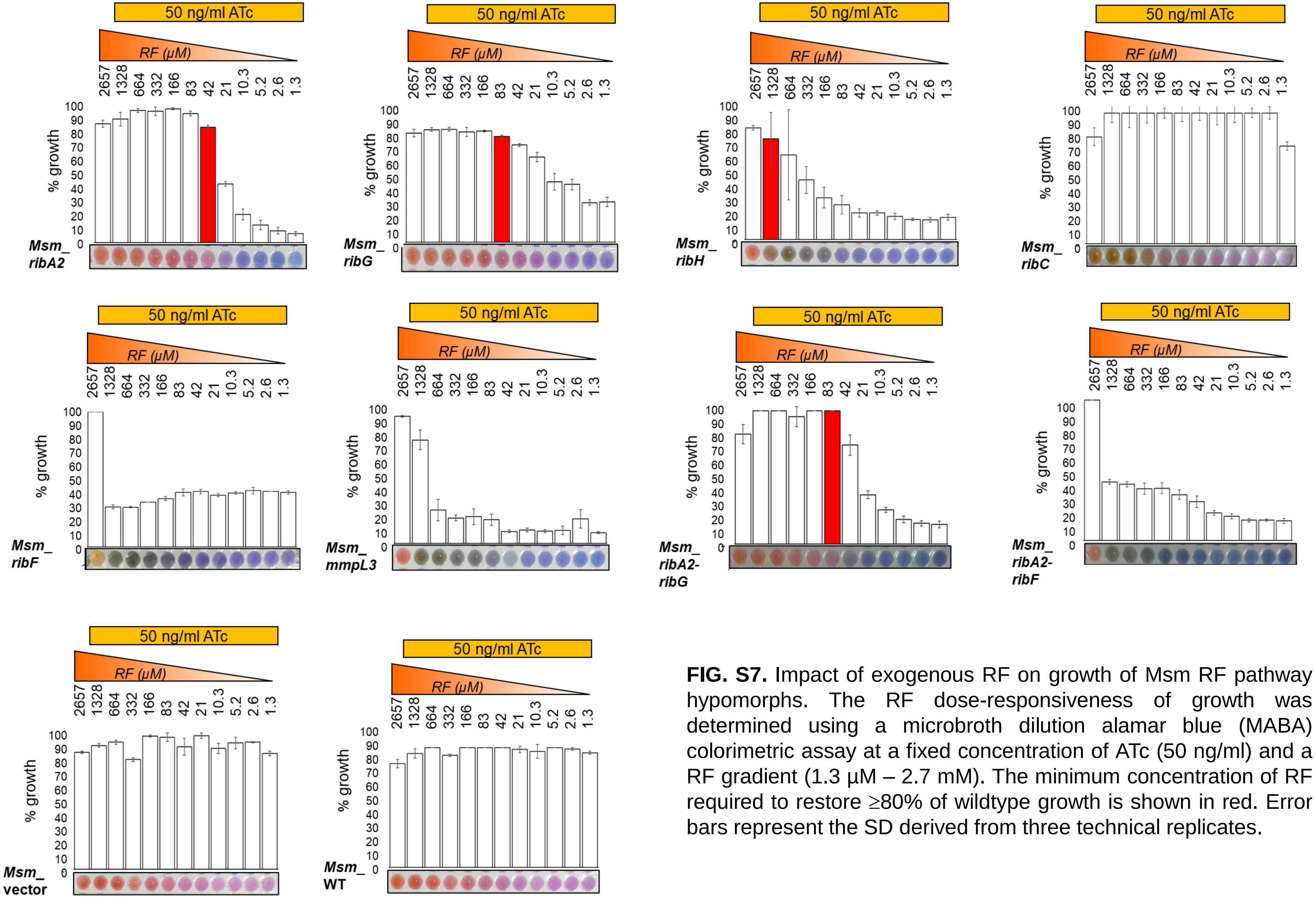

**FIG. S7.** Impact of exogenous RF on growth of Msm RF pathway hypomorphs. The RF dose-responsiveness of growth was determined using a microbroth dilution alamar blue (MABA) colorimetric assay at a fixed concentration of ATc (50 ng/ml) and a RF gradient (1.3 µM – 2.7 mM). The minimum concentration of RF required to restore ≥80% of wildtype growth is shown in red. Error bars represent the SD derived from three technical replicates.

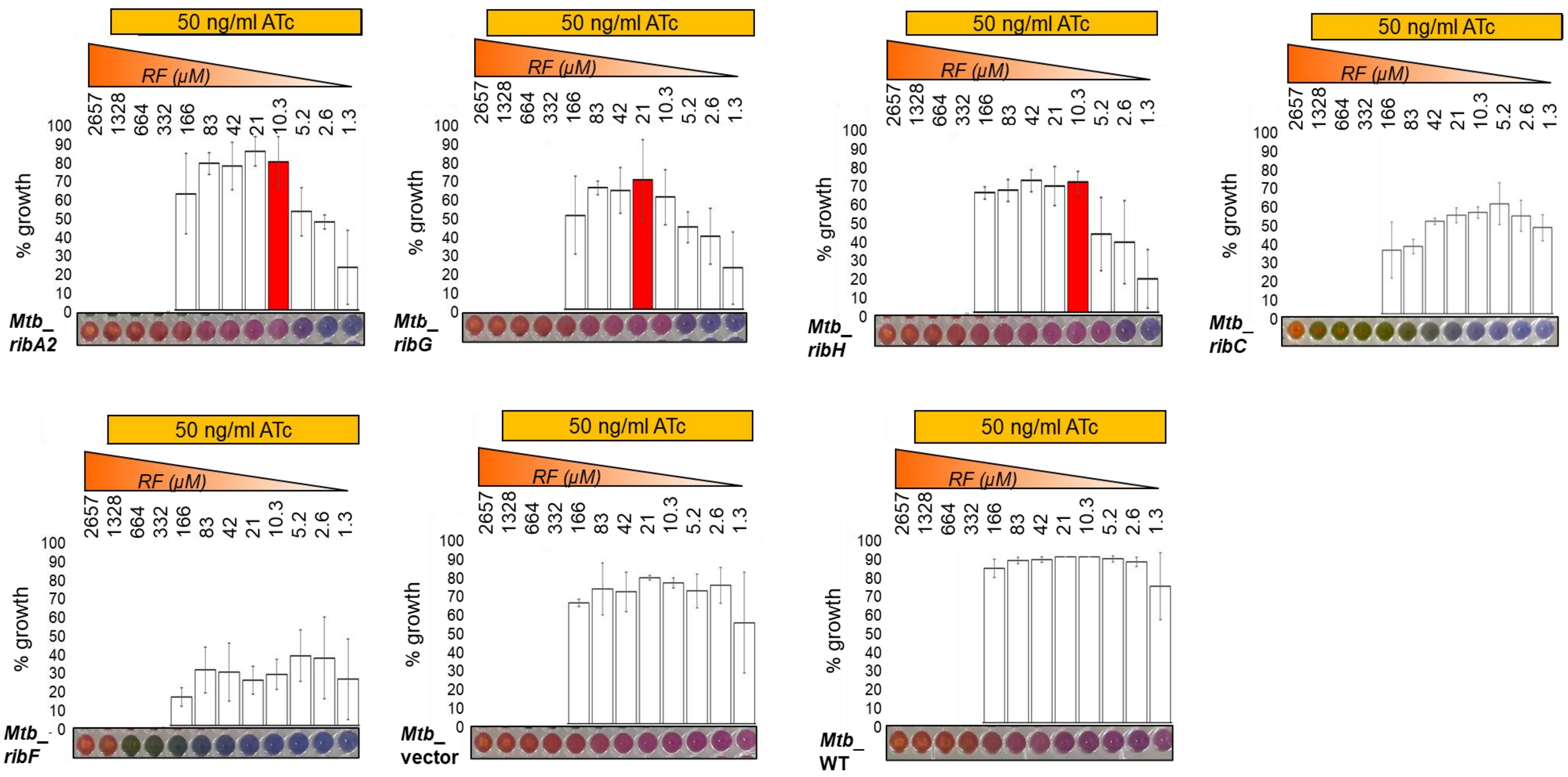

**FIG. S8**. Impact of exogenous RF on growth of Mtb RF pathway hypomorphs. The RF dose-responsiveness of growth was determined using a microbroth dilution alamar blue (MABA) colorimetric assay at a fixed concentration of ATc (50 ng/ml) and a RF gradient (1.3 µM – 2.7 mM). The minimum concentration of RF required to restore growth to ≥80% of the wildtype is shown in red. Error bars represent the SD derived from three technical replicates. Values from the first four wells have been excluded owing to the dense RF pellet obscuring the spectrophotometric reading.

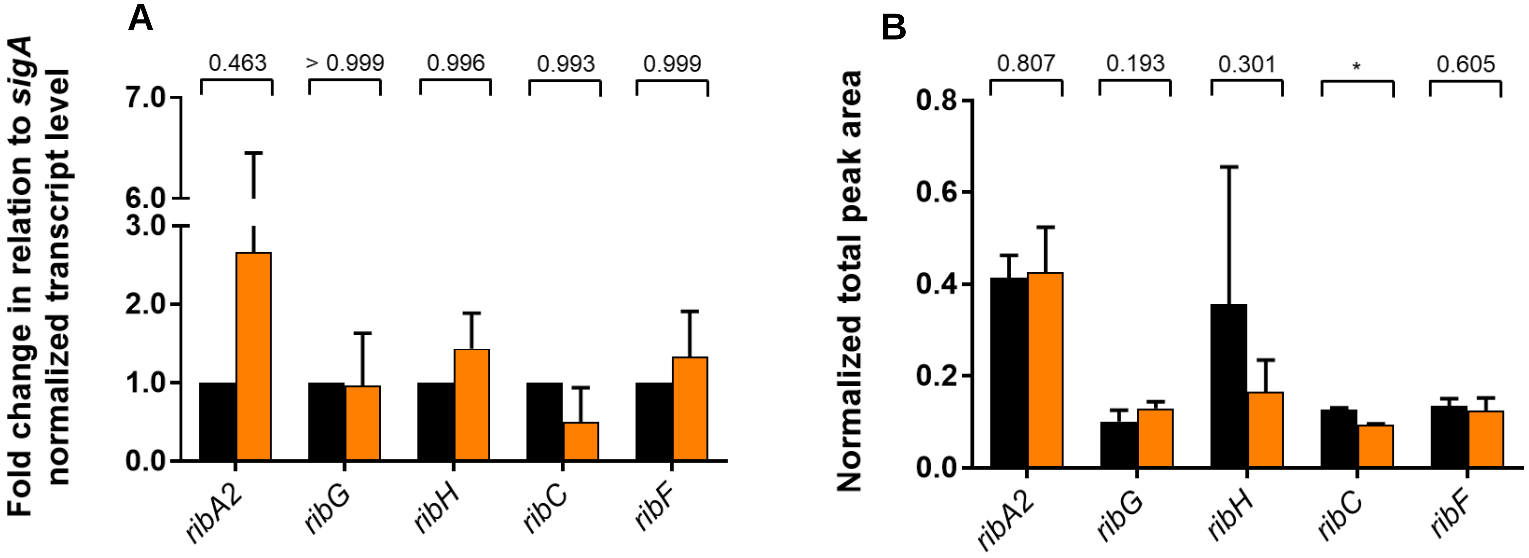

**FIG. S9.** Impact of an exogenous supply of RF on expression of RF pathway genes and abundance of pathway proteins in wildtype Msm. (**A**) The strain was inoculated at $1.5 \times 10^5$-$3 \times 10^5$ CFU/ml and either grown in standard 7H9 media or 7H9 media supplemented with 83 µM RF for 24 hrs prior to harvesting. Fold gene expression in the absence (black) or presence of exogenous RF (orange). Error bars represent SD derived from three biological replicates. Statistical comparisons were performed using a two-way ANOVA and Sidak's multiple comparison test whereby statistical significance is represented by $p < 0.0001$, denoted by an asterisk. (**B**) Relative protein abundance determined by MRM-MS (B) in the absence (black) or presence of RF supplement (orange). Error bars represent the SD derived from three biological replicates. Statistical comparisons were performed by unpaired Student's t-test. Statistical significance is represented by $p<0.05$ and denoted with an asterisk.

**TABLE S1.** Genes involved in RF biosynthesis and utilization in Mtb and Msm

| **Gene** | **Mtb name (H37Rv)** | **Msm name ($mc^2155$)** | **Gene product** | ***In vitro* essentiality in Mtb** | **Vulnerability index (1)[a]** |
|---|---|---|---|---|---|
| *ribA2* | Rv1415 | MSMEG_3072 (*ribAB*) | GTP cyclohydrolase II + 3,4-dihydroxy-2-butanone 4-phosphate synthase (DHBP synthase) | Essential (2-4). Non-essential (5)[b] | -11.8510 (Mtb) -6.9540 (Msm) |
| *ribA1* | Rv1940 | MSMEG_3471 | GTP cyclohydrolase II | Non-essential (2-5) | 1.0210 (Mtb) 0.8120 (Msm) |
| *ribG* | Rv1409 | MSMEG_3067 (*ribD*) | Riboflavin-specific deaminase + 5-amino-6-(5-phosphoribosylamino) uracil reductase | Essential (2-4) Non-essential (5) | -4.8530 (Mtb) -0.2550 (Msm) |
| *ribH* | Rv1416 | MSMEG_3073 MSMEG_6598 (*ribE*) | Probable riboflavin synthase beta chain (lumazine synthase) | Essential (2-4) | -6.4400 (Mtb) -1.9430 (Msm) 0.2170 (Msm) |
| *ribC* | Rv1412 | MSMEG_3071 (*ribE*) | Probable riboflavin synthase alpha chain | Non-essential (2-4) | -5.1780 (Mtb) -1.0910 (Msm) |
| *ribF* | Rv2686c | MSMEG_2653 (*ribF*) | Flavokinase + FAD synthetase | Essential (2-5) | -9.8770 (Mtb) -5.4490 (Msm) |

a. Taken from https://pebble.rockefeller.edu/
b. From Minato *et al.* (ref. 5), essentiality assessed in the presence of riboflavin supplement.

**TABLE S2**. sgRNAs synthesized to create Msm RF pathway hypomorphs.

| Primer name | Primer sequence (5'-3') | Targeted gene |
|---|---|---|
| MSMEG3072_CrispF1 | GGGAGTCCTGGTTCACCGCGTACA | MSMEG_ 3072 (*ribA2*) |
| MSMEG3072_CrispR1 | AAACTGTACGCGGTGAACCAGGAC | |
| MSMEG3072_CrispF2 | GGGAGCCACCGTCCTTGGCGCGCA | |
| MSMEG3072_CrispR2 | AAACTGCGCGCCAAGGACGGTGGC | |
| MSMEG3072_CrispF3 | GGGAGTCGACGGCGGCCTCGGTGTGG | |
| MSMEG3072_CrispR3 | AAACCCACACCGAGGCCGCCGTCGAC | |
| MSMEG3072_CrispF4 | GGGAGCGGAACTCGCCGTGCCGCG | |
| MSMEG3072_CrispR4 | AAACCGCGGCACGGCGAGTTCCGC | |
| MSMEG3072_CrispF5 | GGGAGAACCGAAGACGTCACCCGT | |
| MSMEG3072_CrispR5 | AAACACGGGTGACGTCTTCGGTTC | |
| MSMEG3072_CrispF6 | GGGAGCCCTCGTGTCCGCGCATGT | |
| MSMEG3072_CrispR6 | AAACACATGCGCGGACACGAGGGC | |
| MSMEG3072_CrispF7 | GGGAGCCGTAGTCGCGGGCGTCGG | |
| MSMEG3072_CrispR7 | AAACCCGACGCCCGCGACTACGGC | |
| MSMEG3072_CrispF8 | GGGAGCCGATGCCGTAGTCGCGGGCG | |
| MSMEG3072_CrispR8 | AAACCGCCCGCGACTACGGCATCGGC | |
| MSMEG3072_CrispF9 | GGGAGTTCTCCTTGTTCGCCCGCACCGGCA | |
| MSMEG3072_CrispF9 | AAACTGCCGGTGCGGGCGAACAAGGAGAAC | |
| MSMEG3072_CrispF10 | GGGAGTTCTCCTTGTTCGCCCGCA | |
| MSMEG3072_CrispR10 | AAACTGCGGGCGAACAAGGAGAAC | |
| MSMEG3072_CrispF11 | GGGAATATCGGCTATCGCCCTCTCG | |
| MSMEG3072_CrispR11 | AAACCGAGAGGGCGATAGCCGATAT | |
| MSMEG3072_CrispF12 | GGGAATGCTCTTGTAGCCCACCGC | |
| MSMEG3072_CrispR12 | AAACGCGGTGGGCTACAAGAGCAT | |
| MSMEG3072_CrispF13 | GGGAATCGACCGGATCCCGAGGTCCA | |
| MSMEG3072_CrispR13 | AAACTGGACCTCGGGATCCGGTCGAT | |
| MSMEG3072_CrispF14 | GGGAGCGCTTGGCCGGGTTGTTGG | |
| MSMEG3072_CrispR14 | AAACCCAACAACCCGGCCAAGCGC | |
| MSMEG3072_CrispF15 | GGGAATTCCGTCGGCCTGCGGTCAC | |
| MSMEG3072_CrispR15 | AAACGTGACCGCAGGCCGACGGAAT | |
| MSMEG3067_CrispF1 | GGGAGCCCGCGGCCACCGGGTTGGGG | MSMEG_ 3067 (*ribG*) |
| MSMEG3067_CrispR1 | AAACCCCCAACCCGGTGGCCGCGGGC | |
| MSMEG3067_CrispF2 | GGGAGTCGCGAATTTCCATGTGACAT | |
| MSMEG3067_CrispR2 | AAACATGTCACATGGAAATTCGCGAC | |
| MSMEG3067_CrispF3 | GGGAACGGTCGGCGAGGGTGCCGT | |
| MSMEG3067_CrispR3 | AAACACGGCACCCTCGCCGACCGT | |
| MSMEG3067_CrispF4 | GGGAGTCGTTGAGCACGTTGGCATC | |
| MSMEG3067_CrispR4 | AAACGATGCCAACGTGCTCAACGAC | |
| MSMEG3067_CrispF5 | GGGAGCGAATCGTCGTTGAGCACG | |
| MSMEG3067_CrispR5 | AAACCGTGCTCAACGACGATTCGC | |
| MSMEG3067_CrispF6 | GGGAACCATGGTGCGCGAATCGTCGT | |
| MSMEG3067_CrispR6 | AAACACGACGATTCGCGCACCATGGT | |
| MSMEG3067_CrispF7 | GGGAGGTGCGATCCGACAGCGCAC | |
| MSMEG3067_CrispR7 | AAACGTGCGCTGTCGGATCGCACC | |
| MSMEG3067_CrispF8 | GGGAGCGGTTCACCACGCCCGCGC | |
| MSMEG3067_CrispR8 | AAACGCGCGGGCGTGGTGAACCGC | |
| MSMEG3067_CrispF9 | GGGAGACCGCGGTGATCGGTCCCC | |
| MSMEG3067_CrispF9 | AAACGGGGACCGATCACCGCGGTC | |

| | | |
|---|---|---|
| MSMEG3067_CrispF10 | GGGAACCGCTGCGCGTGCGCGATGC | |
| MSMEG3067_CrispR10 | AAACGCATCGCGCACGCGCAGCGGT | |
| MSMEG3067_CrispF11 | GGGAATCGAGGATCACCGCTCCGACA | |
| MSMEG3067_CrispR11 | AAACTGTCGGAGCGGTGATCCTCGAT | |
| MSMEG3067_CrispF12 | GGGAGCCTGCGACCTGGCCGTCGCGAT | |
| MSMEG3067_CrispR12 | AAACATCGCGACGGCCAGGTCGCAGGC | |
| MSMEG3067_CrispF13 | GGGAACCCGAGGTGACCTCGATGCC | |
| MSMEG3067_CrispR13 | AAACGGCATCGAGGTCACCTCGGGT | |
| MSMEG3073_CrispF1 | GGGAACGCGTCGATCTCGGGCAGAT | MSMEG_ 3073 (*ribH*) |
| MSMEG3073_CrispR1 | AAACATCTGCCCGAGATCGACGCGT | |
| MSMEG3073_CrispF2 | GGGAGTCAGCGACGACGCGTCGATCT | |
| MSMEG3073_CrispR2 | AAACAGATCGACGCGTCGTCGCTGAC | |
| MSMEG3073_CrispF3 | GGGAGGCGACCTTGCGCGCGCCCT | |
| MSMEG3073_CrispR3 | AAACAGGGCGCGCGCAAGGTCGCC | |
| MSMEG3073_CrispF4 | GGGAGCACCCGCACCACCGTCGGGT | |
| MSMEG3073_CrispR4 | AAACACCCGACGGTGGTGCGGGTGC | |
| MSMEG3073_CrispF5 | GGGAGACCGGGATCTCGATGGCGC | |
| MSMEG3073_CrispR5 | AAACGCGCCATCGAGATCCCGGTC | |
| MSMEG3073_CrispF6 | GGGAGGCCAGAGCCTGCGCCACGA | |
| MSMEG3073_CrispR6 | AAACTCGTGGCGCAGGCTCTGGCC | |
| MSMEG3073_CrispF7 | GGGAACACGGGTCAGGCCCTGCGTG | |
| MSMEG3073_CrispR7 | AAACCACGCAGGGCCTGACCCGTGT | |
| MSMEG3073_CrispF8 | GGGAGCCTGCTCCTCGGTGTTGGTGG | |
| MSMEG3073_CrispR8 | AAACCCACCAACACCGAGGAGCAGGC | |
| MSMEG3073_CrispF9 | GGGAGCCCTTGTCCTCGGTGGAAC | |
| MSMEG3073_CrispR9 | AAACGTTCCACCGAGGACAAGGGC | |
| MSMEG3073_CrispF10 | GGGAGCCTGCGCGCCCTTGTCCTCG | |
| MSMEG3073_CrispR10 | AAACCGAGGACAAGGGCGCGCAGGC | |
| MSMEG3071_CrispF1: | GGGAATCTCAAACCAGTCATCACCAATG | MSMEG_ 3071 (*ribC*) |
| MSMEG3071_CrispR1: | AAACCATTGGTGATGACTGGTTTGAGAT | |
| MSMEG3071_CrispF2: | GGGAGCGTGGTCAGCTCGCGAGTGG | |
| MSMEG3071_CrispR2: | AAACCCACTCGCGAGCTGACCACGC | |
| MSMEG3071_CrispF3: | GGGAATCTCAAACCAGTCATCACCA | |
| MSMEG3071_CrispR3: | AAACTGGTGATGACTGGTTTGAGAT | |
| MSMEG3071_CrispF4: | GGGAACCACGTACCGCGAGAGCGCGG | |
| MSMEG3071_CrispR4: | AAACCCGCGCTCTCGCGGTACGTGGT | |
| MSMEG3071_CrispF5: | GGGAATCGGCGGTGAACGCGCCGT | |
| MSMEG3071_CrispR5: | AAACACGGCGCGTTCACCGCCGAT | |
| MSMEG_2653_CrispF1 | GGGAGCCGTCGAACACCCCGATGG | MSMEG_ 2653 (*ribF*) |
| MSMEG_2653_CrispR1 | AAACCCATCGGGGTGTTCGACGGC | |
| MSMEG_2653_CrispF2 | GGGAGCAGGCCCGGATGTAGGTCGA | |
| MSMEG_2653_CrispR2 | AAACTCGACCTACATCCGGGCCTGC | |
| MSMEG_2653_CrispF3 | GGGAGCCTCGACCGTGCGGGTGCGGCC | |
| MSMEG_2653_CrispR3 | AAACGGCCGCACCCGCACGGTCGAGGC | |
| MSMEG_2653_CrispF4 | GGGAGAAGCGCTCACCGGCCTTGC | |
| MSMEG_2653_CrispR4 | AAACGCAAGGCCGGTGAGCGCTTC | |
| MSMEG_2653_CrispF5 | GGGAAGCTGCGCCGGGTGGTTACC | |
| MSMEG_2653_CrispR5 | AAACGGTAACCACCCGGCGCAGCT | |
| MSMEG_2653_CrispF6 | GGGAGTCGGAGGTGAACGGCATCAC | |
| MSMEG_2653_CrispR6 | AAACGTGATGCCGTTCACCTCCGAC | |

| | | |
|---|---|---|
| MSMEG_2653_CrispF7 | GGGAGGCGGTGCCACGTTGGCCGT | |
| MSMEG_2653_CrispR7 | AAACACGGCCAACGTGGCACCGCC | |
| MSMEG_2653_CrispF8 | GGGAGTCTTGCCCACGCCAGCGTTGC | |
| MSMEG_2653_CrispR8 | AAACGCAACGCTGGCGTGGGCAAGAC | |
| MSMEG_2653_CrispF9 | GGGAGACGACGTGCAGGTGCTCGA | |
| MSMEG_2653_CrispR9 | AAACTCGAGCACCTGCACGTCGTC | |
| MSMEG_2653_CrispF10 | GGGAGTCGGAGGTGAACGGCATCA | |
| MSMEG_2653_CrispR10 | AAACTGATGCCGTTCACCTCCGAC | |
| MSMEG_MmpL3_CrispF1 | GGGAGCGACAGACTGGCTGCCCTCGTC | MSMEG_ 0250 |
| MSMEG_MmpL3_CrispR1 | AAACGACGAGGGCAGCCAGTCTGTCGC | |

**TABLE S3**. sgRNAs synthesized to create Mtb RF pathway hypomorphs.

| **Primer name** | **Primer sequence (5'-3')** | **Targeted gene** |
|---|---|---|
| Rv1415_CrispF1 | GGGAGTCGCAGATGGCACCGTCCA | Rv1415 (*ribA2*) |
| Rv1415_CrispR1 | AAACTGGACGGTGCCATCTGCGAC | |
| Rv1415_CrispF2 | GGGAACCCCACCATCCTTGGCCCGCA | |
| Rv1415_CrispR2 | AAACTGCGGGCCAAGGATGGTGGGGT | |
| Rv1415_CrispF3 | GGGAGCACTCCGAATGCACCCGGA | |
| Rv1415_CrispR3 | AAACTCCGGGTGCATTCGGAGTGC | |
| Rv1415_CrispF4 | GGGAGACCCAAACACATCGCCGGT | |
| Rv1415_CrispR4 | AAACACCGGCGATGTGTTTGGGTC | |
| Rv1415_CrispF5 | GGGAGCCCTCGTGGCCACGCATGT | |
| Rv1415_CrispR5 | AAACACATGCGTGGCCACGAGGGC | |
| Rv1415_CrispF6 | GGGAGCCGATCCCGTAATCCCTTGCG | |
| Rv1415_CrispR6 | AAACCGCAAGGGATTACGGGATCGGC | |
| Rv1415_CrispF7 | GGGAATCGAACGTACCCCAAGATCGA | |
| Rv1415_CrispR7 | AAACTCGATCTTGGGGTACGTTCGAT | |
| Rv1415_CrispF8 | GGGAACCCGCTTGGCCGGGTTGTTGG | |
| Rv1415_CrispR8 | AAACCCAACAACCCGGCCAAGCGGGT | |
| Rv1415_CrispF9 | GGGAGTTCTCCGCGTTGGCCCGCACCGGCA | |
| Rv1415_CrispR9 | AAACTGCCGGTGCGGGCCAACGCGGAGAAC | |
| Rv1415_CrispF10 | GGGAACAAGGCACCGCCGAATTCTC | |
| Rv1415_CrispR10 | AAACGAGAATTCGGCGGTGCCTTGT | |
| Rv1415_CrispF12: | GGGAGCCTCGGTGTGGCCGGGCCGGC | |
| Rv1415_CrispR12: | AAACGCCGGCCCGGCCACACCGAGGC | |
| Rv1415_CrispF13: | GGGAGCTTGCGCCGCCATTCGATCAAG | |
| Rv1415_CrispR13: | AAACCTTGATCGAATGGCGGCGCAAGC | |
| Rv1415_CrispF14: | GGGAGCGAAACTCCCCATGACGAG | |
| Rv1415_CrispR14: | AAACCTCGTCATGGGGAGTTTCGC | |
| Rv1409_CrispR1 | AAACCCGACGACCCGGCCCTGACC | Rv1409 (*ribG*) |
| Rv1409_CrispF2 | GGGAATCATGGTGCGTGCCTCGTCGT | |
| Rv1409_CrispR2 | AAACACGACGAGGCACGCACCATGAT | |
| Rv1409_CrispF3 | GGGAGGTGCGATCCGACAACGCCC | |
| Rv1409_CrispR3 | AAACGGGCGTTGTCGGATCGCACC | |
| Rv1409_CrispF4 | GGGAGAGGGTGGGACCTCCTTCCA | |
| Rv1409_CrispR4 | AAACTGGAAGGAGGTCCCACCCTC | |
| Rv1409_CrispF5 | GGGAGGCGAGGGTGGGACCTCCTT | |
| Rv1409_CrispR5 | AAACAAGGAGGTCCCACCCTCGCC | |
| Rv1409_CrispF6 | GGGAATCCGGTTGATCGCACCCGCTC | |
| Rv1409_CrispR6 | AAACGAGCGGGTGCGATCAACCGGAT | |
| Rv1409_CrispF7 | GGGAACCGCGGTAACCGGACCGCCCA | |
| Rv1409_CrispR7 | AAACTGGGCGGTCCGGTTACCGCGGT | |
| Rv1409_CrispF8 | GGGAGCCCGGCGCAGCGCCACCACC | |
| Rv1409_CrispR8 | AAACGGTGGTGGCGCTGCGCCGGGC | |
| Rv1409_CrispF9 | GGGAATGGTGACCACCACGATGGCG | |
| Rv1409_CrispR9 | AAACCGCCATCGTGGTGGTCACCAT | |
| Rv1409_CrispF10 | GGGAGGCGTACTTCCAGGTGACAT | |
| Rv1409_CrispR10 | AAACATGTCACCTGGAAGTACGCC | |
| Rv1409_CrispF11: | GGGAATCCGGTTGATCGCACCCGCTCG | |
| Rv1409_CrispR11: | AAACCGAGCGGGTGCGATCAACCGGAT | |
| Rv1409_CrispF12: | GGGAATAAGTCGTGCCTTTGACCTG | |

| | | |
|---|---|---|
| Rv1409_CrispR12: | AAACCAGGTCAAAGGCACGACTTAT | |
| Rv1409_CrispF13: | GGGAGCCGGCGCCGACGATCCGACCG | |
| Rv1409_CrispR13: | AAACCGGTCGGATCGTCGGCGCCGGC | |
| Rv1416_CrispF1: | GGGAACGCATCCAGCGACGGCAGAT | Rv1416 (*ribH*) |
| Rv1416_CrispR1: | AAACATCTGCCGTCGCTGGATGCGT | |
| Rv1416_CrispF2: | GGGAGCACACCAGACGCATCCAGCG | |
| Rv1416_CrispR2: | AAACCGCTGGATGCGTCTGGTGTGC | |
| Rv1416_CrispF3: | GGGAACCACCGGAATCTCGATCGCGC | |
| Rv1416_CrispR3: | AAACGCGCGATCGAGATTCCGGTGGT | |
| Rv1416_CrispF4: | GGGAGGCCAATTCCTGCGCCACCA | |
| Rv1416_CrispR4: | AAACTGGTGGCGCAGGAATTGGCC | |
| Rv1416_CrispF5: | GGGAATCACGACGCCAAGTGCGACG | |
| Rv1416_CrispR5: | AAACCGTCGCACTTGGCGTCGTGAT | |
| Rv1412_CrispF1: | GGGAGCGCCGAGCCCGGAGACCGTC | Rv1412 (*ribC*) |
| Rv1412_CrispR1: | AAACGACGGTCTCCGGGCTCGGCGC | |
| Rv1412_CrispF2: | GGGAGCGCCGAGCCCGGAGACCGTC | |
| Rv1412_CrispR2: | AAACGACGGTCTCCGGGCTCGGCGC | |
| Rv1412_CrispF3: | GGGAGCGTGGTCAGCTCCCGGGTCG | |
| Rv1412_CrispR3: | AAACCGACCCGGGAGCTGACCACGC | |
| Rv1412_CrispF4: | GGGAGCACCACTTCCCAGTGCTCG | |
| Rv1412_CrispR4: | AAACCGAGCACTGGGAAGTGGTGC | |
| Rv1412_CrispF5: | GGGAGTCGGCGGTGAATTGGCCGT | |
| Rv1412_CrispR5: | AAACACGGCCAATTCACCGCCGAC | |
| Rv2786c_CrispF1 | GGGAGCTTCGACGGTGCGGGTGCGTCC | Rv2786c (*ribF*) |
| Rv2786c_CrispR1 | AAACGGACGCACCCGCACCGTCGAAGC | |
| Rv2786c_CrispF2 | GGGAGCAGGACCGGATGTAGGTGGA | |
| Rv2786c_CrispR2 | AAACTCCACCTACATCCGGTCCTGC | |
| Rv2786c_CrispF3 | GGGAATCGGCGGCGCCACGTTCGCGG | |
| Rv2786c_CrispR3 | AAACCCGCGAACGTGGCGCCGCCGAT | |
| Rv2786c_CrispF4 | GGGAACGCAGGACCGGATGTAGGT | |
| Rv2786c_CrispR4 | AAACACCTACATCCGGTCCTGCGT | |
| Rv2786c_CrispF5 | GGGAATCGGTGGTGAACGGCATCAC | |
| Rv2786c_CrispR5 | AAACGTGATGCCGTTCACCACCGAT | |
| Rv2786c_CrispF6 | GGGAAGCGGCGTACACGCCGTCGG | |
| Rv2786c_CrispR6 | AAACCCGACGGCGTGTACGCCGCT | |
| Rv2786c_CrispF7 | GGGAACCTCCACCACATGTAGGTGCTCGA | |
| Rv2786c_CrispR7 | AAACTCGAGCACCTACATGTGGTGGAGGT | |
| Rv2786c_CrispF8 | GGGAATCGGTGGTGAACGGCATCA | |
| Rv2786c_CrispR8 | AAACTGATGCCGTTCACCACCGAT | |
| Rv2786c_CrispF9 | GGGAACCATGTCGCCGGCGTCCAC | |
| Rv2786c_CrispR9 | AAACGTGGACGCCGGCGACATGGT | |
| Rv2786c_CrispF10 | GGGAGTGCGGGTGCGTCCGGAGAAG | |
| Rv2786c_CrispR10 | AAACCTTCTCCGGACGCACCCGCAC | |
| Mtb_MmpL3_CrispF1: | GGGAACCCGGTTAACCAGCTTGCC | Rv0206c (*mmpL3*) |
| Mtb_MmpL3_CrispR1: | AAACGGCAAGCTGGTTAACCGGGT | |

**TABLE S4.** ORBIT oligo to create targeted knockout in Msm *ribH*.

| Primer name | Primer | Application |
|---|---|---|
| MSMRibH_ORB_KO | TGCACGTCCCAGGTCTCGTCCGCCATCTGTGA GGCCATCGTCACGCGCGGCGCAATTCGCGCAG CGTCACGGTTTGTCTGGTCAACCACCGCGGTC TCAGTGGTGTACGGTACAAACCGGGCAGATCG GGGACGCCGACGCCGCTCACTGCGCCCCGCCC GATTCCGTCGGCCTGCGGTCACCCAGCA | Msm *ribH* knockout |

**TABLE S5.** Plasmids used in this study.

| Plasmid | Description | Reference |
|---|---|---|
| pLJR962 | ATc-inducible ($P_{tet}$) *Streptococcus thermophilius* (Sth) dCas9 and $P_{tet}$ sgRNA, integrating at the L5 locus in *Msm*; $Kan^R$ | (6) |
| pLJR965 | ATc-inducible ($P_{tet}$) Sth dCas9 and $P_{tet}$ sgRNA, integrating at the L5 locus in *Mtb*; $Kan^R$ | (6) |
| pKM461 | Plasmid expresses $P_{tet}$ Che9c phage RecT annealase and $P_{tet}$ Bxb1 phage integrase; *oriE, oriM, sacB* ; $Kan^R$, $Tet^R$ | (7) |
| pKM464 | Non-replicating vector carrying *attB*; $Hyg^R$ | (7) |
| pLJR962_*ribA2* | Derivative of pLJR962 carrying a *ribA2* (MSMEG_3072) targeting sgRNA; $Kan^R$ | This study |
| pLJR962_*ribG* | Derivative of pLJR962 carrying a *ribG* (MSMEG_3067) targeting sgRNA; $Kan^R$ | This study |
| pLJR962_*ribH* | Derivative of pLJR962 carrying a *ribH* (MSMEG_3073) targeting sgRNA; $Kan^R$ | This study |
| pLJR962_*ribC* | Derivative of pLJR962 carrying a *ribC* (MSMEG_3071) targeting sgRNA; $Kan^R$ | This study |
| pLJR962_*ribF* | Derivative of pLJR962 carrying a *ribF* (MSMEG_2653) targeting sgRNA; $Kan^R$ | This study |
| pLJR962_*ribA2_ribG* | Derivative of pLJR962 carrying *ribA2* (MSMEG_3072) and *ribG* (MSMEG_3067) targeting sgRNAs; $Kan^R$ | This study |
| pLJR962_*ribA2_ribF* | Derivative of pLJR962 carrying a *ribA2* (MSMEG_3072) and a *ribF* (MSMEG_2653) targeting sgRNAs; $Kan^R$ | This study |
| pLJR962_*mmpL3* | Derivative of pLJR962 carrying a *mmpL3* (MSMEG_0250) targeting sgRNA; $Kan^R$ | This study |
| pLJR962_6598 | Derivative of pLJR962 carrying a MSMEG_6598 targeting sgRNA; $Kan^R$ | This study |
| pLJR965_*ribA2* | Derivative of pLJR965 carrying a *ribA2* (Rv1415) targeting sgRNA | This study |
| pLJR965_*ribG* | Derivative of pLJR965 carrying a *ribG* (Rv1409) targeting sgRNA | This study |
| pLJR965_*ribH* | Derivative of pLJR965 carrying a *ribH* (Rv1416) targeting sgRNA | This study |
| pLJR965_*ribC* | Derivative of pLJR965 carrying a *ribC* (Rv1412) targeting sgRNA | This study |
| pLJR965_*ribF* | Derivative of pLJR965 carrying a *ribF* (Rv2786c) targeting sgRNA | This study |

| | | |
|---|---|---|
| pLJR965_*mmpL3* | Derivative of pLJR965 carrying a *mmpL3* (Rv0206c) targeting sgRNA | This study |

**TABLE S6.** Strains used in this study.

| Strain | Description | Reference |
|---|---|---|
| *Escherichia coli* DH5α | $F^{-}$ φ80*lac*ZΔM15 Δ(*lac*ZYA-*arg*F)U169 *rec*A1 *end*A1 *hsd*R17($r_K^-$, $m_K^+$) *pho*A *sup*E44 $\lambda^-$ *thiA*-1 *gyr*A96 *rel*A1 | New England Biolabs |
| $mc^2155$ | High frequency transformation mutant of *M. smegmatis* $mc^26$ ATCC®™ | (8) |
| *Msm_ribA2* | Anhydrotetracycline (ATc)-inducible Msm hypomorph targeting *ribA2* generated using plasmid pLJR962_*ribA2* | This study |
| *Msm_ribG* | ATc-inducible Msm hypomorph targeting *ribG* generated using plasmid pLJR962_*ribG* | This study |
| *Msm_ribH* | ATc-inducible Msm hypomorph targeting *ribH* generated using plasmid pLJR962_*ribH* | This study |
| *Msm_ribC* | ATc-inducible Msm hypomorph targeting *ribC* generated using plasmid pLJR962_*ribC* | This study |
| *Msm_ribF* | ATc-inducible Msm hypomorph targeting *ribF* generated using plasmid pLJR962_*ribF* | This study |
| *Msm_ribA2_ribG* | ATc-inducible Msm hypomorph targeting *ribA2* and *ribG* generated using plasmid pLJR962_*ribA2_ribG* | This study |
| *Msm_ribA2_ribF* | ATc-inducible Msm hypomorph targeting *ribA2* and *ribF* generated using plasmid pLJR962_*ribA2_ribF* | This study |
| *Msm_mmpL3* | ATc-inducible Msm hypomorph targeting *mmpL3* generated using plasmid pLJR962_*mmpL3* | This study |
| *Msm*_vector | ATc-inducible Msm control strain generated using pLJR962 and no targeting sgRNA | This study |
| *Msm*_MSMEG_6598 | ATc-inducible Msm hypomorph targeting MSMEG_6598 generated using plasmid pLJR962_ MSMEG_6598 | This study |
| *Msm* Δ*ribH* | In-frame deletion mutant lacking internal region in *ribH* generated using construct pKM464 and directed oligo; $Hyg^R$ marked mutant generated using ORBIT. This strain is not a RF auxotroph. | This study |
| *Msm* Δ*ribH* MSMEG_6598 | ATc-inducible Msm hypomorph targeting MSMEG_6598 generated using plasmid pLJR962_ MSMEG_6598 in *Msm* Δ*ribH* | This study |
| H37RvMA | Mtb H37Rv isolate ATCC® 27294™ virulent laboratory strain | (9) |
| *Mtb_ribA2* | ATc-inducible Mtb hypomorph targeting *ribA2* generated using plasmid pLJR965_*ribA2* | This study |
| *Mtb_ribG* | ATc-inducible Mtb hypomorph targeting *ribG* generated using plasmid pLJR965_*ribG* | This study |
| *Mtb_ribH* | ATc-inducible Mtb hypomorph targeting *ribH* generated using plasmid pLJR965_*ribH* | This study |
| *Mtb_ribC* | ATc-inducible Mtb hypomorph targeting *ribC* generated using plasmid pLJR965_*ribC* | This study |
| *Mtb_ribF* | ATc-inducible Mtb hypomorph targeting *ribF* generated using plasmid pLJR965_*ribF* | This study |
| *Mtb*_vector | ATc-inducible Mtb strain carrying pLJR965 and no targeting sgRNA | This study |

| | | |
|---|---|---|
| *Mtb_mmpL3* | ATc-inducible Mtb strain carrying pLJR965 targeting *mmpL3* generated using plasmid pLJR965 *mmpL3* | This study |

**TABLE S7.** Primers used for qRT-PCR analyses.

| **Primer name** | **Primer sequence (5'-3')** | **Amplicon size (bp)** | **Gene region amplified** |
|---|---|---|---|
| qRT_MSM_3072Fwd | GCCGACGATTTCACCAAGC | 159 | MSMEG_3072 (*ribA2*) |
| qRT_MSM_3072Rev | GTCCTTCTGGCTGACGATCTC | | |
| qRT_MSM_3067Fwd | TCAACCGTTGCGTGTCGT | 151 | MSMEG_3067 (*ribG*) |
| qRT_MSM_3067Rev | GCCCTCGAGCATGATGTC | | |
| qRT_MSM_3073Fwd | CACCGAGATCTGCGACGC | 102 | MSMEG_3073 (*ribH*) |
| qRT_MSM_3073Rev | TGTCTCACCTTGGATCACGAC | | |
| qRT_MSM_3071Fwd | TTGGTGATGACTGGTTTGAGA | 106 | MSMEG_3071 (*ribC*) |
| qRT_MSM_3071Rev | ATGACGTCCACTTCCAGGTTC | | |
| qRT_MSM_2653Fwd | GATCGACGTGTTCCTGGTGAT | 163 | MSMEG_2653 (*ribF*) |
| qRT_MSM_2653Rev | CAGCATGTCGACGTTGCC | | |
| qRT_MSM_0250Fwd | AACGATCCGGAGAAGATGTGG | 128 | MSMEG_0250 (*mmpL3*) |
| qRT_MSM_0250Rev | CGCAACTCGTCGATCTTCTTG | | |
| qRT_MSM_2758Fwd | ACCAAGGGCTACAAGTTCTCG | 198 | MSMEG_2758 (*sigA*) |
| qRT_MSM_2758Rev | CATCTCCTTGGCGAGCTCTTC | | |
| qRT_Rv1415Fwd | CACGGAATGGCATTGGAACTG | 144 | Rv1415 (*ribA2*) |
| qRT_Rv1415Rev | GCAGAACCCCACCATCCTTG | | |
| qRT_Rv1409Fwd | CATGTCACCTGGAAGTACGCC | 104 | Rv1409 (*ribG*) |
| qRT_Rv1409Rev | CGATGCAGATCCAGGCGT | | |
| qRT_Rv1416Fwd | AATTGGCCCGCAATCATGATG | 130 | Rv1416 (*ribH*) |
| qRT_Rv1416Rev | GGCGTCGAGGAATCCAGC | | |
| qRT_Rv1412Fwd | ACATCGTGCAGGGACATGTG | 123 | Rv1412 (*ribC*) |
| qRT_Rv1412Rev | AGCCCTTTTCGACGACATAGC | | |
| qRT_Rv2786Fwd | GCTTGGTTCACGGTGCTC | 103 | Rv2786c (*ribF*) |
| qRT_Rv2786Rev | CGGAGAAGGTGGGATTGGTC | | |
| qRT_Rv0206Fwd | GACCACCACGATCGTCTTGAT | 129 | Rv0206c (*mmpL3*) |
| qRT_Rv0206Rev | TGTCCGTCGACGAATATCCAC | | |
| qRT_Rv2703Fwd | CGGTGATTTCGTCTGGGATGA | 113 | Rv2703 (*sigA*) |
| qRT_Rv2703Rev | TGCCGATCTGTTTGAGGTAGG | | |

**TABLE S8.** Primers used to amplify junctions between adjacent genes to interrogate operonic structure.

| **Primer name** | **Primer sequence (5'-3')** | **Amplicon size (bp)** | **Region amplified** |
|---|---|---|---|
| Msm_ribC_ribA2Fwd | GGGTACGGGTCGGTAAGAG | 166 | Junction between *ribC* and *ribA2* in Msm |
| Msm_ribC_ribA2Rev | CGCCCTCTCGACGGAATC | | |
| Msm_ribA2_ribHFwd | CGGTGAGCATGGACGACT | 110 | Junction between *ribA2* and *ribH* in Msm |
| Msm_ribA2_ribHRev | TCTCGGGCAGATCGGGGA | | |
| Mtb_ribC_Rv1413Fwd | CGTAGTCGCAAAGTATGTTGAGC | 331 | Junction between *ribC* and Rv1413 in Mtb |
| Mtb_ribC_Rv1413Rev | CAATGGGTAGGTGATGAATACGT | | |
| Mtb_Rv1413_Rv1414Fwd | CTTCCCGATCAGCTGCCG | 155 | Junction between Rv1413 and Rv1414 in Mtb |
| Mtb_Rv1413_Rv1414Rev | AACGGTCAGCGCGATGTC | | |
| Mtb_Rv1414_ribA2Fwd | GCAACCCTGATTGATCGCTG | 170 | Junction between Rv1414 and *ribA2* in Mtb |
| Mtb_Rv1414_ribA2Rev | GTCCAACCTCGTCATCTTTGC | | |
| Mtb_ribA2_ribHFwd | GACAAATTGGGGCACGACTTG | 172 | Junction between *ribA2* and *ribH* in Mtb |
| Mtb_ribA2_ribHRev | ATCTTTCCGTGCCAGCTG | | |

**TABLE S9.** Peptides used for targeted proteomics by MRM-MS

| Protein[¥] | Peptide (m/z) | Transitions | Retention time (min) | LOD (nM) | LOQ (nM) |
|---|---|---|---|---|---|
| RibA2 | R.LGLLPMYAVNQD**K**.H<br>(731.3921 ++) | P [y9] - 1065.5034+<br>Y [y7] - 837.4101+<br>A [y6] - 674.3468+<br>V [y5] - 603.3097+<br>N [y4] - 504.2413+ | 7.2 | 45.97 | 139.3 |
| | R.GEISGPGSDGDDVLV**R**.V*<br>(786.8786++) | P [y11] - 1129.5484+<br>S [y9] - 975.4742+ | 5.4 | 6.35 | 19.24 |

| | | | | | |
|---|---|---|---|---|---|
| | | D [y8] - 888.4421+<br>G [y7] - 773.4152+<br>D [y5] - 601.3668+ | | | |
| RibG | R.VVYAVADPNPVAAGGSA**R**.M*<br>(857.4496++) | P [y11] - 996.5221+<br>P [y9] - 785.4264+<br>A [y6] - 518.2681+<br>G [y5] - 447.2310+<br>G [y4] - 390.2096+ | 5.4 | 7.81 | 23.67 |
| | R.EVSSDANVLNDDS**R**.T | A [y9] - 1003.4803+ | 6.4 | 4.03 | 12.22 |

| | | | | | |
|---|---|---|---|---|---|
| | (760.8448 ++) | N [y8] - 932.4432+<br>V [y7] - 818.4003+<br>L [y6] - 719.3319+<br>N [y5] - 606.2478+ | | | |
| RibH | K.VAADAGIPDPTVV**R**.V<br>(690.8777++) | D [y11] - 1139.6055+<br>A [y10] - 1024.5786+<br>G [y9] - 953.5415+<br>I [y8] - 896.5200+<br>P [y7] - 783.4359 | 5.7 | 5.4 | 16.38 |

| | | | | | |
|---|---|---|---|---|---|
| | R.VLGAIEIPVVAQALA**R**.T<br><br>(810.4958++) | I [y12] - 1279.7732+<br><br>E [y11] - 1166.6892+<br><br>I [y10] - 1037.6466+<br><br>P [y9] - 924.5625+<br><br>V [y7] - 728.4413+ | 8.7 | 26.62 | 80.67 |
| RibC | R.SSLAGVAVGD**R**.V<br><br>(516.2776++) | L [y9] - 857.4839+<br><br>A [y8] - 744.3999+<br><br>G [y7] - 673.3628+<br><br>V [y6] - 616.3413+ | 4.8 | 4.49 | 13.59 |

| | | | | | |
|---|---|---|---|---|---|
| | | A [y5] - 517.2729+ | | | |
| | R.IALPPALS**R**.Y*<br><br>(469.2951++) | A [y8] - 824.4989+<br><br>L [y7] - 753.4617+<br><br>P [y6] - 640.3777+<br><br>P [y5] - 543.3249+ | 6.1 | 5.42 | 16.43 |
| RibF | R.YIHELLVEHLHVVEVVVGENFTFG**K**.K<br><br>(969.8516+++) | G [y8] - 450.2165++<br><br>N [y6] - 357.1845++<br><br>F [y5] - 300.1630++<br><br>T [y4] - 226.6288++ | N/A | N/A | N/A |

| | | | | | |
|---|---|---|---|---|---|
| | | F [y3] - 176.1050++ | | | |
| | R.NETVTFSSTYI**R**.A<br><br>(l709.3515++) | V [y9] - 1073.5626+<br><br>T [y8] - 974.4942+<br><br>F [y7] - 873.4465+<br><br>S [y6] - 726.3781+ | 5.9 | 5.01 | 15.19 |

¥Mtb protein names used. RibF peptide 'YIH' was not able to be validated with a labeled synthetic peptide, so it was not included for sample testing.
